# Ecological network analysis of watershed meta-ecosystems: A new perspective on quantifying the integrated watershed ecosystem

**DOI:** 10.1101/2021.05.28.446148

**Authors:** Haile Yang

**Affiliations:** Institute of Biodiversity Science at Fudan University, Shanghai, 200438, China; Department of Ecology and Evolutionary Biology, School of Life Sciences, Fudan University, Shanghai, 200438, China

**Keywords:** Drainage basin, catchment, watershed hierarchy system, subwatershed, spatial process, integrated watershed ecosystem evolution

## Abstract

A watershed is an integrated ecosystem. In different disciplines, a watershed has been described as a geomorphic unit, a hydrological unit, an ecological unit, or a socio-economic unit and has been quantitatively described using different indicator systems. Until now, no general indicator system has existed that could quantitatively describe the geomorphic features, hydrologic features, ecological features and socio-economic features of an integrated watershed ecosystem (IWE) simultaneously. Here, we proposed a quantitative descriptive framework for an IWE (QDFIWE). This QDFIWE involved three steps: (1) constructing a watershed meta-ecosystem (WME) based on the hierarchical system of the watershed; (2) constructing flow networks based on the WME; and (3) identifying the holistic properties (such as spatial throughput, spatial organization and spatial resilience) of the WME through analyzing its flow networks based on ecological network analysis (ENA). Then, we applied this method to study the geomorphic topological structure, geomorphic spatial structure, natural water resource system and integrated water resource system of the Yangtze River basin. The results suggested that based on the QDFIWE, (1) one could construct different WMEs and corresponding flow networks for different requirements; (2) one could obtain time series of the holistic properties of an IWE to analyze its evolution; (3) one could compare, classify and cluster any number of IWEs through identifying their holistic properties according to similar strategies; and (4) one could determine or create more indicators, which could provide additional information, based on the holistic properties of an IWE. This study demonstrates that the QDFIWE is a general method of quantifying the holistic properties of all subsystems of an IWE simultaneously. Thus, the method can improve the understanding of the IWE.

## 1 Introduction

In ecology, a watershed had been regarded as a basic ecosystem (Lotspeich, 1980). However, more precisely, a watershed is an integrated ecosystem (Thurow and Juo, 1995; Kolok et al., 2009; Cabello et al., 2015). Historically, in different disciplines, watersheds have been described as geomorphic units (Horton, 1945; Strahler, 1957; Ozdemir and Bird, 2009), hydrologic units (Seaber et al., 1987; Huang and Ferng, 1990), ecological units (Lotspeich, 1980; Frissell et al., 1986; Hornbeck and Swank, 1992), and socio-economic units (Wang and Zhang, 2014) quantitatively described by different indicators. As watersheds have become the spatially based unit of the integrated governance of watershed, models that integrate water, ecosystems, social systems, economic systems, health and well-being have become increasingly proposed. These models include the “Patuxent landscape model”, “watershed governance prism”, and “complex holarchic social-ecological system” (Costanza et al., 2002; Parkes et al., 2010; Cabello et al., 2015). However, until now, no general indicator system could quantitatively describe the geomorphic subsystem, hydrologic subsystem, ecological subsystem and socio-economic subsystem of an integrated watershed ecosystem (IWE) simultaneously.

It is necessary to develop a quantitative framework to completely and thoroughly describe the IWE if we want to accurately understand how IWEs function. Because the hierarchical structure of a watershed is a common feature of the geomorphic subsystem (Ozdemir and Bird, 2009), hydrologic subsystem (Seaber et al., 1987), ecological subsystem (Frissell et al., 1986) and socio-economic subsystem (Wang and Zhang, 2014), we believe that information-based ecological network analysis (ENA) indicators can be used to quantify the spatial throughput, spatial organization and spatial resilience of an IWE and indicate the corresponding spatial growth, spatial development and spatial stabilization. Based on ENA of an IWE, the spatially holistic properties of each subsystem in an IWE can be identified, and these properties can be combined to describe the overall holistic properties of the IWE, such as to assess the sustainable development of an ecological economic system in a watershed.

ENA is a system-oriented method used to identify holistic properties, especially those not evident through direct observation, by focusing on the interactions among the components of the system (Ulanowicz, 1986; Patricio et al., 2004; Huang and Ulanowicz, 2014). From a flow network perspective, growth usually implies an increase and/or expansion, which may involve either a greater spatial extent or the increase in a flow medium. This growth is commonly quantified as any increase in total system throughput (TST). Development, however, implies an increase in organization, which is independent of growth and can be represented by an increase in the average mutual information (AMI) inherent in a network. The combined actions of growth and development, as they pertain to increases in network size and organization, can be represented as an increase in system ascendency (A). The mathematical upper bound of system ascendency is called the development capacity (C), which is measured by the diversity of system flows and normalized by the TST. Redundancy (R) is generated by structural ambiguities derived from a variety of system inputs, exports, dissipations and internal exchanges (functional redundancy). R quantifies the residual “freedom” of a system and represents its potential for recovery or innovative restructuring.

Traditionally, ENA addresses the relationships between the individual components and the overall properties of a system by analyzing material–energy–information networks (Ulanowicz, 1997; Goerner et al., 2009; Huang and Ulanowicz, 2014). Here, to quantify the IWE, similar concepts and network methods must be created to describe the spatial structure and spatial processes of the IWE. Then, TST, AMI and R are used to quantify the spatial throughput, spatial organization and spatial resilience of the IWE, respectively.

The objectives of this study are threefold: (1) to propose a quantitative descriptive framework for an IWE (QDFIWE) based on an ENA of the WME; (2) to provide a case study (quantifying the holistic properties of the geomorphic topological structure, geomorphic spatial structure, natural water resource system and integrated water resource system of the Yangtze River basin) that illustrates how to quantify an IWE based on the QDFIWE; and (3) to describe the spatial throughput, spatial organization and spatial resilience of the Yangtze River basin based on the geomorphic topological structure, geomorphic spatial structure, natural water resource system and integrated water resource system.

## 2 Methodology

QDFIWE is mainly used to quantify the spatial throughput, spatial organization and spatial resilience of an IWE. This framework involves three steps: (1) constructing a watershed meta-ecosystem (WME) based on the hierarchical system of the watershed; (2) constructing flow networks based on the WME; and (3) identifying the spatial throughput, spatial organization and spatial resilience of the WME through analyzing the flow networks based on ENA.

### 2.1 Construction of the WME

The basic assumption of this study is as follows: a WME is constructed of subwatersheds (SWs) and corresponding main stream regions (MSRs). The delineation of SWs and corresponding MSRs involves three steps: (1) set the minimum area of an SW; (2) select the strategy of assigning stream order; and (3) delineate the SWs and MSRs.

“Setting the minimum area of an SW” means “setting the boundary conditions of initial streams”. The strategies of assigning stream order can be classified into three basic types: (1) binary tree strategies, (2) weighted binary tree strategies, and (3) main-branch strategies. Based on the stream orders, the catchment area of every initial channel, which is larger than the minimum area of an SW, is delineated as an SW. Then, the catchment area of the following channel of a definite order is delineated as an MSR.

In the binary tree strategy, three stream-ordering methods exist. The first is Shreve’s stream magnitude (Figure 1a). This ordering method assigns a value of one to every initial channel. The magnitude of the following channel is the sum of magnitudes of its tributaries. Thus, the magnitude of a particular link is the number of initial streams which contribute to it. The second method is Scheidegger’s stream magnitude (Figure 1b). This ordering method assigns a value of two to every initial channel. The magnitude of the following channel is the sum of the magnitudes of its tributaries minus one. Thus, the magnitude of a particular link is the number of initial streams that contribute to it plus one. The third method is topological dimension stream ordering (Figure 1c). This method assigns a value of one to the main stream at the watershed outlet. Then, a value of one is added for each contributing channel as long as bifurcation occurs. Based on these three stream-ordering strategies, every stream segment has a defined order that differs from those of adjacent stream segments. Additionally, each segment with a definite order is associated with a catchment area, i.e., an SW or MSR.

**Figure 1.**
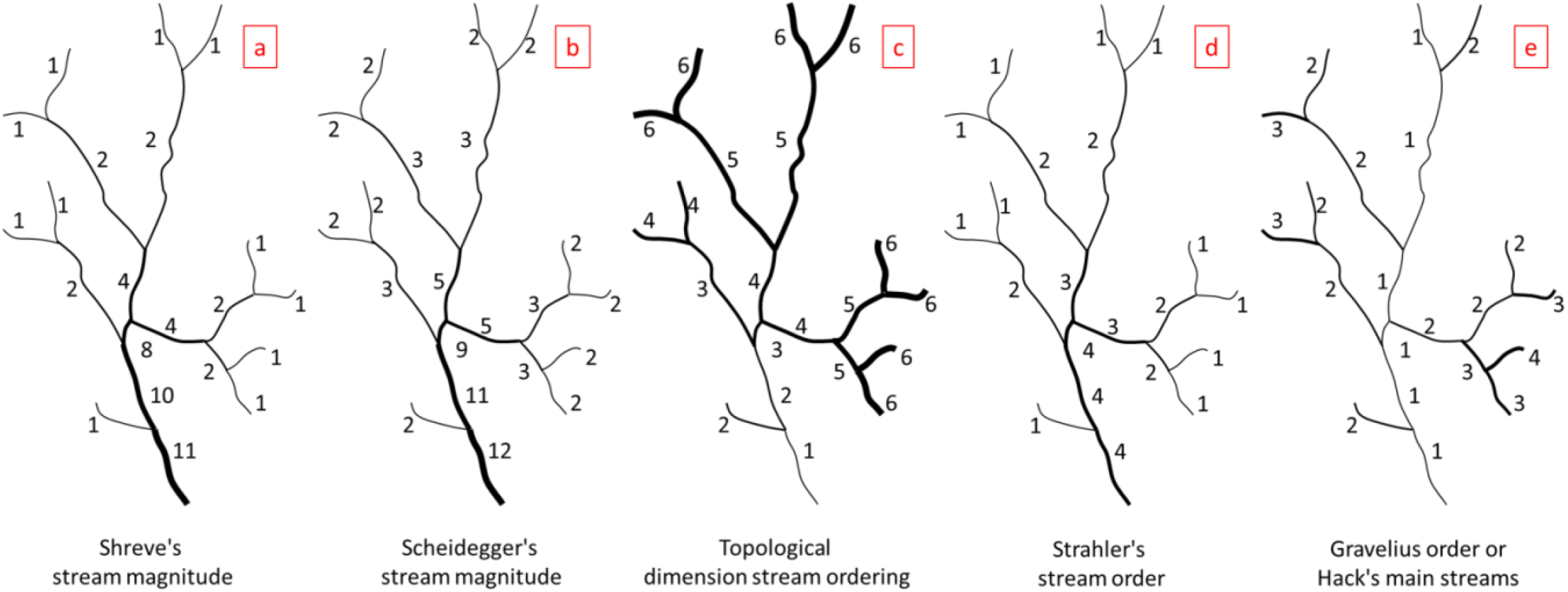
Schematic representation of stream-ordering principles based on binary tree (a, b, c), weighted binary tree (d) and main-branch (e) strategies

The main weighted binary tree strategy is the stream-ordering method proposed by Strahler (Figure 1d). In Strahler’s stream order, the order of the initial channel is assigned a value of one. If a channel with one and only one tributary has the largest Strahler order i and all other tributaries have orders less than i, then the order of the stream remains i. However, if a channel has two or more tributaries with greatest order i, then its order get i+1. Based on this ordering strategy, some adjacent stream segments have the same order. However, every segment with a definite order has a specific catchment area, i.e., an SW or MSR. Moreover, adjacent segments of the same order have a common catchment area, i.e., a continuous MSR.

The most popular main-branch strategies are the Gravelius order and Hack’s main stream methods (Figure 1e). These methods are the same. Thus, in this approach, the main stream is set to one, and, consequently, all its tributaries are assigned an order of two. Then, their tributaries are assigned an order of three. This process continues until all streams are ordered. The order of every stream remains constant up to its initial link, and the route of every main stream is determined according to the maximum flow length of a particular stream. Obviously, the watershed of the main stream (i.e., the stream with order one) is the watershed itself, and it can be divided into a set of SWs (i.e., the catchment areas of streams of order two) and an MSR (i.e., the remainder of the catchment area of the order one stream minus the catchment area of order two streams). Of course, the area of each SW must be larger than the minimum area of an SW.

In practice, some integrated strategies can be used to delineate SWs and MSRs, such as the delineation of hydrologic units by the United State Geological Survey (USGS) and the delineation of watershed resource divisions by the Ministry of Water Resources of the People’s Republic of China (MWR).

If the SWs and corresponding MSRs of a watershed are described by a directed graph, the WME can be constructed.

### 2.2 Construction of flow networks

The basic assumption of flow network construction is as follows: the flow network that represents the system of interest can be constructed from diverse perspectives based on traditional flows or spatial structures. In a WME, one could construct flow networks based on geomorphic features (such as the topological structure, spatial structure, etc.), hydrologic features (such as the hydrologic cycle, sediment transport, biogeochemical cycle, etc.), ecological features (i.e., ecological spatial structure/processes driven by the hydrologic cycle), or socio-economic features (i.e., the socio-economic spatial structure/processes related to the watershed spatial structure).

The flow network of a geomorphic topological structure can always be described using a directed graph of a WME (Figure 2a). In addition, the flow network of a geomorphic spatial structure can always be described using a weighted directed graph of a WME in which every node (i.e., SW or MSR) has a weight (such as the catchment area, drainage density, basin shape and so on) (Figure 2b). The catchment area always relates to total runoff; the drainage density always relates to the erosion degree and sediment transport; and the basin shape always relates to the features of the flood peak.

**Figure 2.**
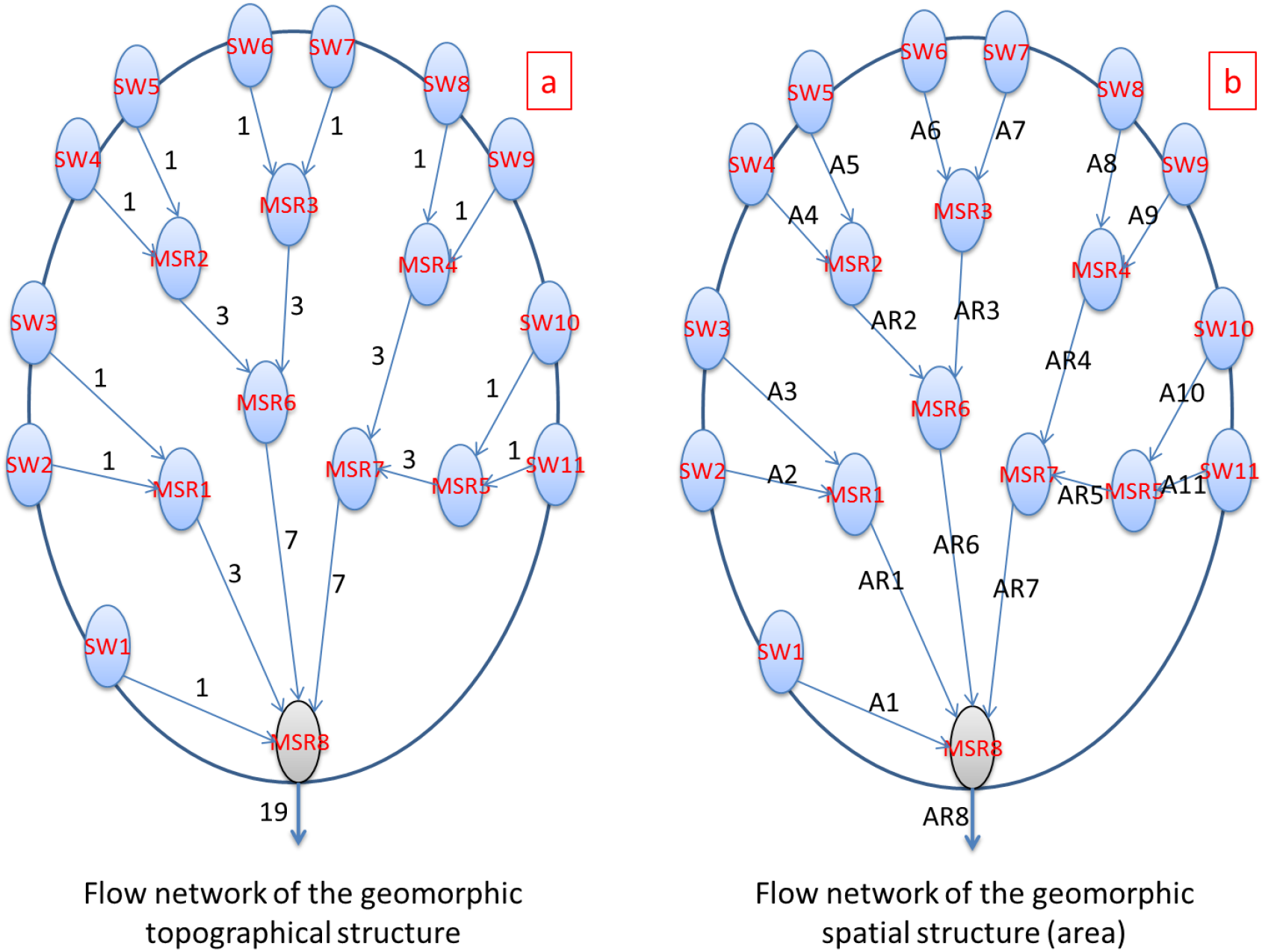
Schematic representations of flow networks of the geomorphic topological structure (a) and geomorphic spatial structure (b) constructed based on Strahler’s stream order. SW: subwatershed; MSR: main stream region; A1: the area of SW1; and AR1: the area of MSR1 and its upstream catchment area (SW2 and SW3).

The flow network of the hydrologic cycle can always be described using a weighted directed graph of a WME in which every node (i.e., SW or MSR) has a 4-element structure (i.e., precipitation, evapotranspiration, input and output) (Figure 3a). In the flow network of sediment transport, every node also has a 4-element structure (i.e., erosion, deposition, input and output) (Figure 3b). Moreover, in the flow network of the biogeochemical cycle, every node has a 4-element structure (i.e., leaching, absorption, input and output) (Figure 3c). The hydrologic cycle always relates to natural water resources; sediment transport always relates to geomorphic evolution; and the biogeochemical cycle always relates to environmental and ecological dynamics.

**Figure 3.**
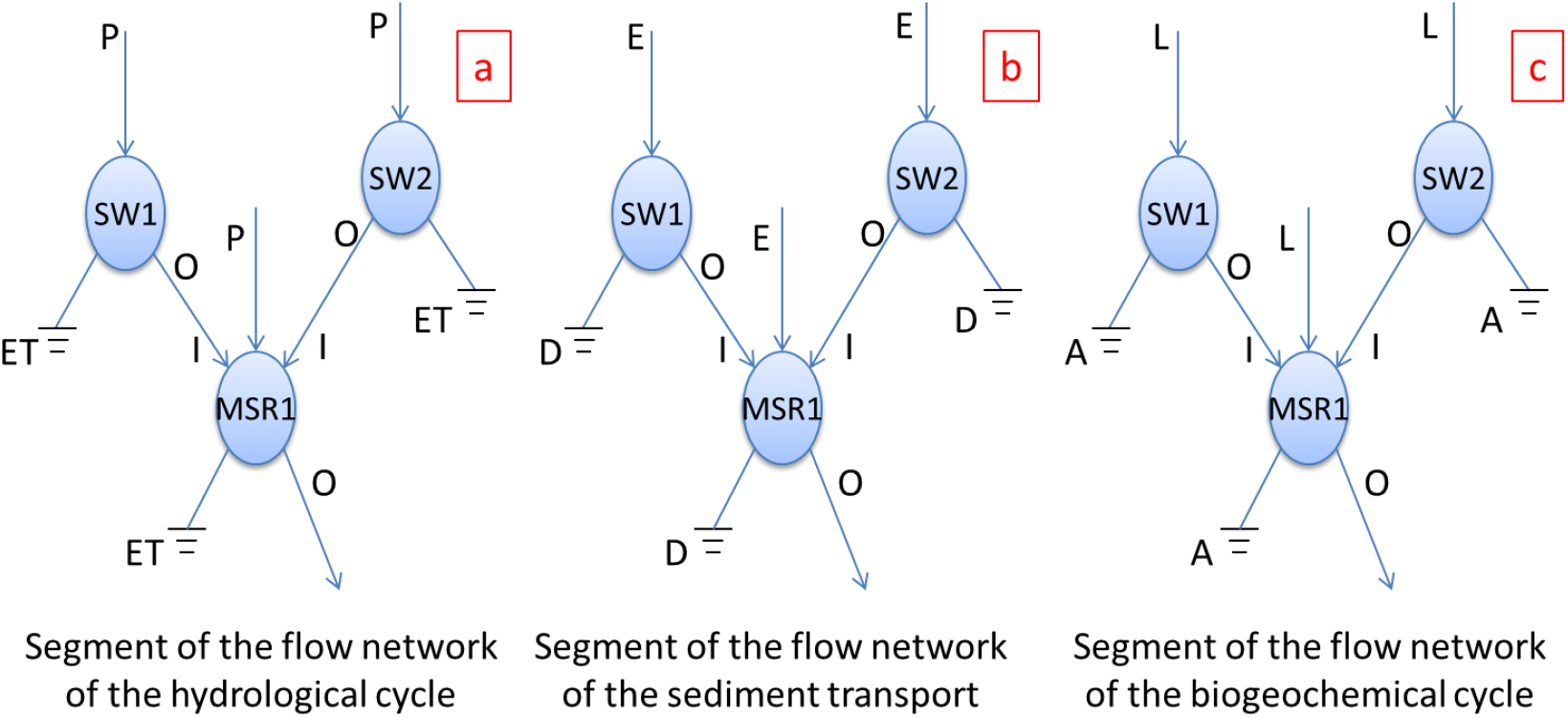
Schematic representations of the flow network segments of the hydrologic cycle (a), sediment transport (b) and biogeochemical cycle (c) with 4-element structures. SW: subwatershed; MSR: main stream region; I: input; O: output; P: precipitation; ET: evapotranspiration; E: erosion; D: deposition; L: leaching; and A: absorption.

The flow network of the ecological spatial structure driven by hydrologic processes can always be described using a weighted directed graph of a WME in which every node (i.e., SW or MSR) has a weight (such as habitat area for a specific species or community). Additionally, the flow network of ecological spatial processes driven by the hydrologic cycle can always be described using a weighted directed graph of a WME in which every node (i.e., SW or MSR) has a 5-element structure (such as gross primary production, respiration, accumulation, input and output for ecosystem production or subsidy, respiration, feedback, input and output for land-water interactions; Figure 4a-b), even has a multi-element structure (such as food web with inlet and outlet along the drainage system) (Figure 4c).

**Figure 4.**
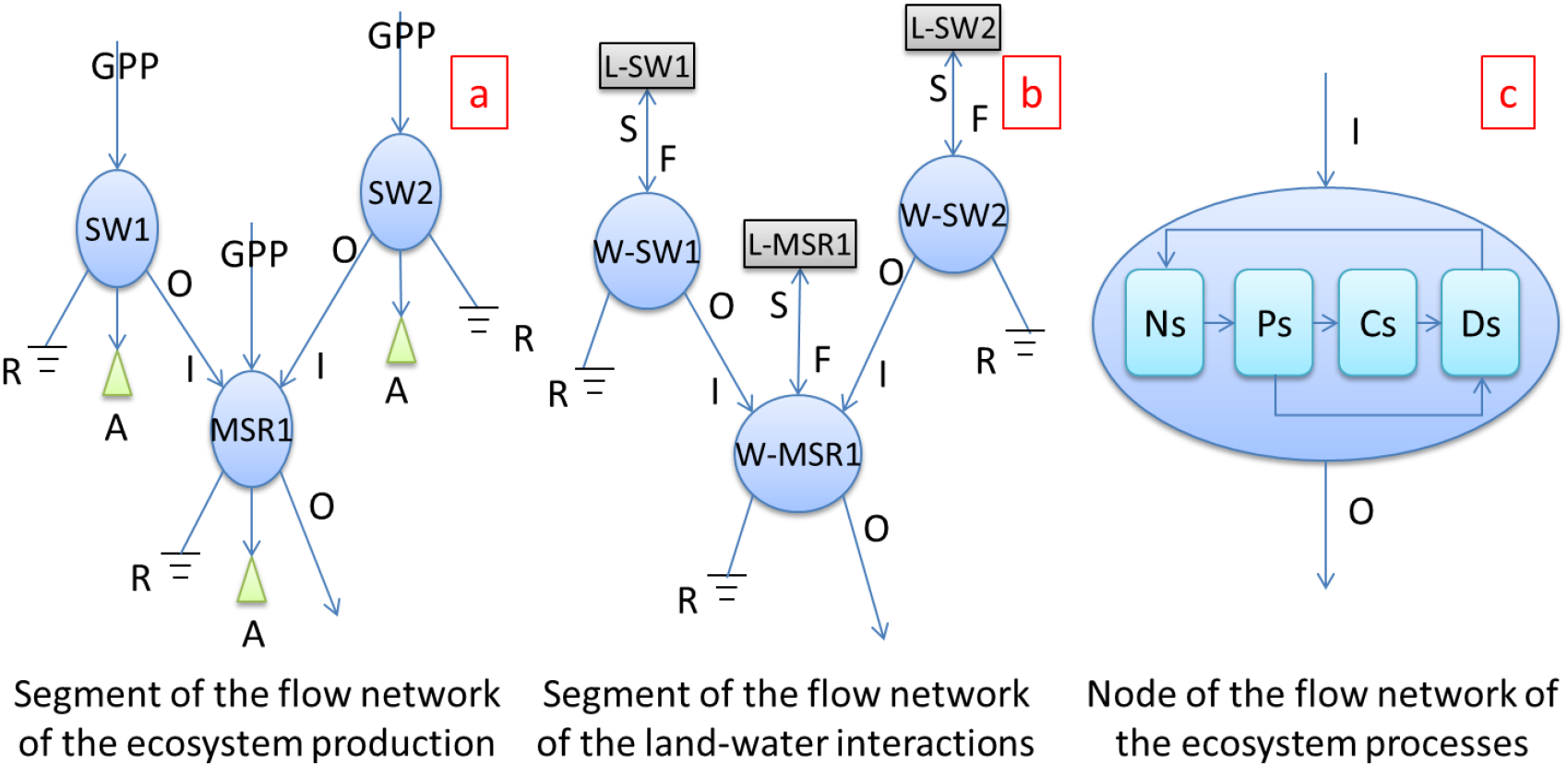
Schematic representations of the flow network segments of ecological interactions with 5-element structures (a, b) and of the flow network node of ecosystem processes with 6-element structures (c). SW: subwatershed; MSR: main stream region; L: land; W, water; I: input; O: output; GPP: gross primary production; R: respiration; A: accumulation; S: subsidy; F: feedback; Ns: inorganic nutrients; Ps: producers; Cs: consumers; Ds: decomposers.

The flow network of the socio-economic spatial structure related to the watershed spatial structure can always be described using a weighted directed graph of a WME in which every node (i.e., SW or MSR) has a weight (such as gross domestic product (GDP), population, the number of core cities and so on). The flow network of socio-economic spatial processes related to the watershed spatial structure can always be described using a weighted directed graph of a WME in which every node (i.e., SW or MSR) has an integrated structure (such as the water use system combined with the natural water resource system, sewage discharge combined with ecosystem services, production-transportation-consumption combined with the associated resource and environmental footprints, population mobility combined with wealth flows and so on) (Figure 5).

**Figure 5.**
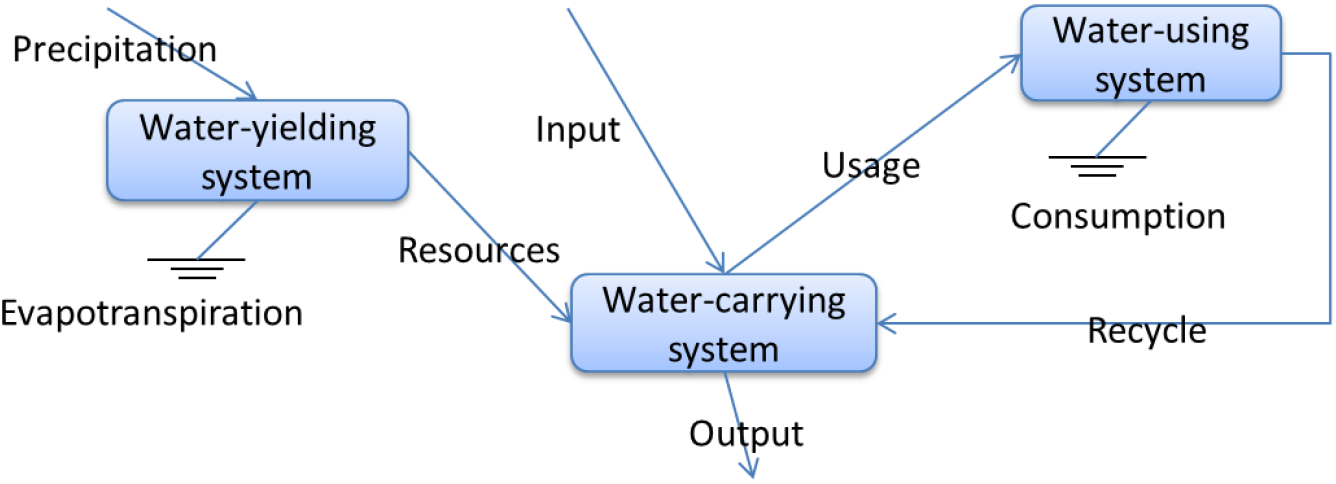
Schematic representation of the flow network node of an integrated water resource system with 3-compartments

The flow network of a watershed spatial structure with multiple features can be variable and described using a weighted directed graph of a WME in which every node (i.e., SW or MSR) has an integrated structure.

### 2.3 ENA of a flow network

Based on the flow network of a WME, the holistic properties (TST, C, H, A, AMI, R, and SR (defined below)) can be identified by ENA. TST is simply the sum of all flows in a meta-ecosystem, and it reflects the size and overall activity of the meta-ecosystem (Ulanowicz, 1986; Patricio et al., 2004; Huang and Ulanowicz, 2014):

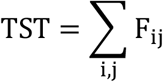

where *F_ij_* denotes the flow of a medium from node *i* to node *j*.

AMI is an ecological information-based index used to estimate the organization of a meta-ecosystem, and H serves as an upper bound of the AMI (Ulanowicz, 1986; Patricio et al., 2004; Huang and Ulanowicz, 2014):

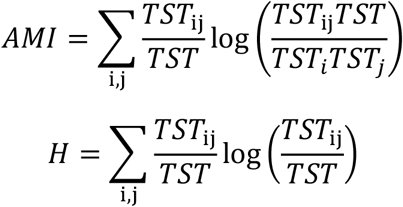

where *TST*_*i*_ = ∑_*j*_ *F*_ij_, *TST*_*j*_ = ∑_*i*_ *F*_ij_ and *H* ≥ *AMI* ≥ 0.

Ascendency (A) is a key property of a network of flows that quantifies both the level of system activity and the degree of organization (Ulanowicz, 1986; Patricio et al., 2004; Huang and Ulanowicz, 2014).

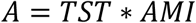

C is the diversity of the system flows scaled by the TST. Additionally, C is as an upper bound of system ascendency (Ulanowicz, 1986; Patricio et al., 2004; Huang and Ulanowicz, 2014).

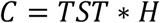

R is the degree to which pathways parallel each other in a network, which can be regarded as resilience, an attribute that is complementary (opposite) to ascendency (Ulanowicz, 1986; Patricio et al., 2004; Huang and Ulanowicz, 2014).

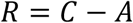

Specific redundancy (SR) measures the total flexibility of the system on a per-unit-flow basis. It consists mostly of pathway redundancy and the varieties of external inputs and outputs in open systems (Ulanowicz, 1986; Patricio et al., 2004; Huang and Ulanowicz, 2014).

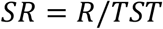

Traditionally, statistical analyses of topographic watershed characteristics (such as channel profiles, drainage areas, catchment slopes and so on) have provided many useful clues to geomorphic evolution (Kirby and Whipple, 2001; Miller et al., 2013). Now, an ENA of the geomorphic properties of a WME could provide a new approach to analyzing geomorphic evolution.

Watershed resilience, the representative capacity of a watershed to absorb and recover from perturbations or disturbances, has been traditionally analyzed based on water quality time series (Hoque et al., 2012; Hoque et al., 2016), water quantity time series (Qi et al., 2016), multiple hydrologic features (Hazbavi and Sadeghi, 2016; Sadeghi and Hazbavi, 2017) or multiple socio-ecological properties (Nemec et al., 2014). Here, the ENA of a WME provides a new perspective for the quantification of watershed resilience.

Above all, the ENA of a WME provides a general framework for analyzing, comparing, classifying and clustering any number of IWEs based on their variable flow networks.

### 2.4 Study area

With a length of 6,300 km, the Yangtze River is the third longest river in the world. The Yangtze Basin, which extends for some 3,200 km from west to the east and for more than 1,000 km from north to south, drains an area of 1,808,500 km^2^. The geomorphic features of the basin are very complex, and the evolution of its fluvial systems has been the focus of numerous studies (Clark et al., 2004; Zheng et al., 2013; Zheng, 2015).

The Yangtze River basin is comparatively rich in water resources. The average yearly rainfall amounts to approximately 1,100 mm, with high spatio-temporal variation. Therefore, the spatial and temporal variabilities in precipitation (Zhang et al., 2008; Chen et al., 2014), the runoff and suspended sediment load changes (Zhang et al., 2006a; Zhang et al., 2006b; Yang et al., 2015; Zhao et al., 2015), and the complex river-lake interactions (Hu et al., 2007; Lai et al., 2013; Zhang et al., 2015) have been studied in detail.

The Yangtze River basin is the main part of the gigantic Yangtze River Economic Belt „super zone’ of China. In the Yangtze River basin in 2015, the population was 454.56 million; water usage reached 205.46 billion cubic meters; and the GDP increased to ¥ 24.46 trillion (http://www.cjw.gov.cn/zwzc/bmgb/). Thus, flood disasters (Yin and Li, 2001; Liu et al., 2014), water usage (Chen et al., 2001; Okadera et al., 2014), water pollution (Chen et al., 2000; Qi et al., 2014), the deterioration and conservation of biodiversity (Fu et al., 2003; Fang et al., 2006; Huang and Li, 2016), and the human impacts on nutrient flux variations (Li et al., 2007; Wang et al., 2014; Chen et al., 2016) have been studied previously.

### 2.5 Data sources

To quantify the holistic properties of the geomorphic topological structure, geomorphic spatial structure, natural water resource system and integrated water resource system of the Yangtze River basin, data sets of geomorphic features, hydrologic features and water usage in the Yangtze River basin are needed. Series of these data are available from the Changjiang & Southwest Rivers Water Resources Bulletin provided by the Changjiang Water Resources Commission of the MWR (http://www.cjw.gov.cn/zwzc/bmgb/).

## 3 Results

### 3.1 Construction of the Yangtze WME

Theoretically, to construct the Yangtze WME, the SWs and corresponding MSRs should be delineated based on a defined strategy. However, because a well-established division system (i.e., watershed resource divisions by the Chinese MWR) exists and all data are collected based on this system, here, the Yangtze WME was constructed based on the watershed resource divisions of the Yangtze River basin (Figure 6) (http://www.cjw.gov.cn/zwzc/bmgb/).

**Figure 6.**
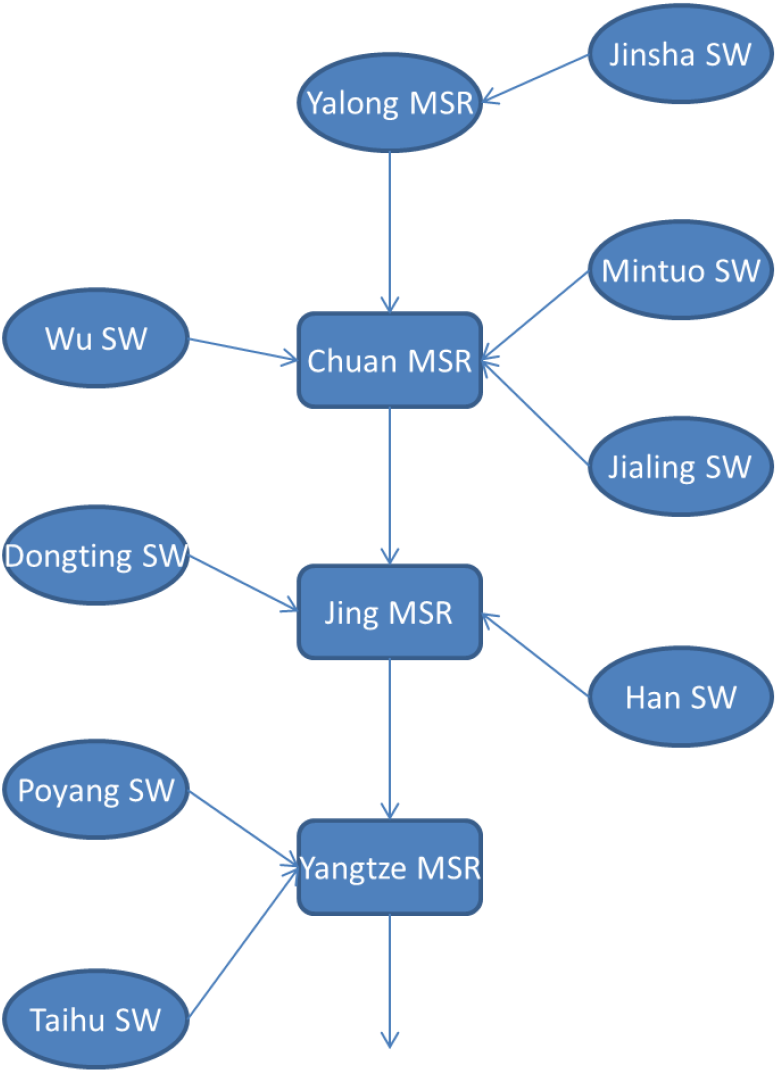
Schematic diagram of the Yangtze watershed meta-ecosystem (WME) constructed based on the watershed resource divisions of the Yangtze River basin (http://www.cjw.gov.cn/zwzc/bmgb/). SW: subwatershed; MSR: main stream region; Jinsha SW: the catchment of Jinsha upper Shigu; Yalong MSR: the catchment of Jinsha from Shigu to Yibin, including the catchment of Yalong; Chuan MSR: the main stream catchment from Yibin to Yichang; Jing MSR: the main stream catchment from Yichang to Hukou; Yangtze MSR: the main stream catchment from Hukou to estuary.

### 3.2 Construction of flow networks

Based on the Yangtze WME, four flow networks that represented the geomorphic topological structure (Figure 7a), geomorphic spatial structure (Figure 7b), natural water resource system (Figure 8), and integrated water resource system (Figure 9, a node of the flow network) of the Yangtze River basin were constructed.

**Figure 7.**
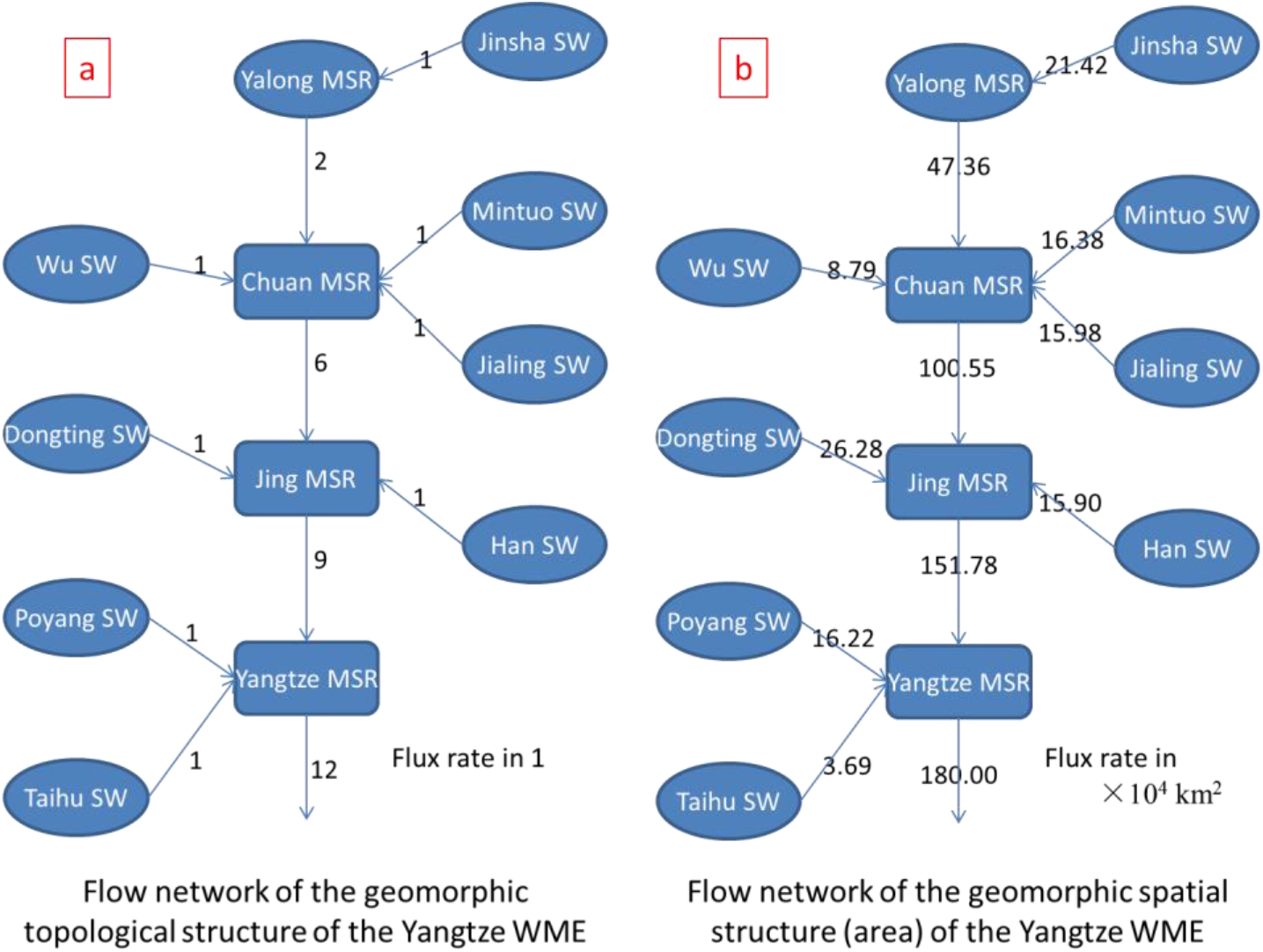
Schematic diagrams of the flow networks of the geomorphic structures of the Yangtze watershed meta-ecosystem (WME). SW: subwatershed and MSR: main stream region.

**Figure 8.**
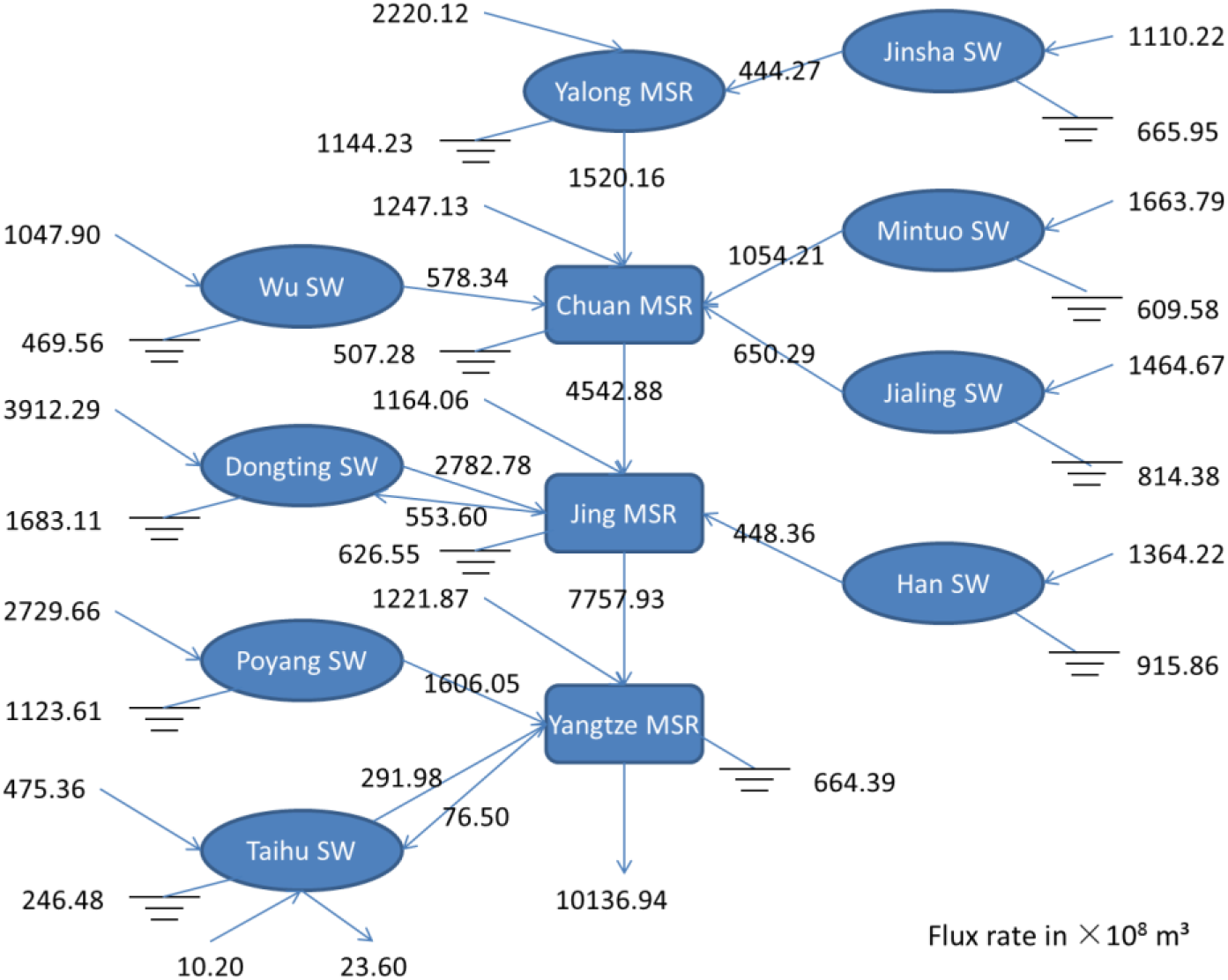
Schematic diagram of the flow network of the natural water resource system (with precipitation, evapotranspiration, input and output) in the Yangtze watershed meta-ecosystem (WME) (2014). SW: subwatershed and MSR: main stream region.

**Figure 9.**
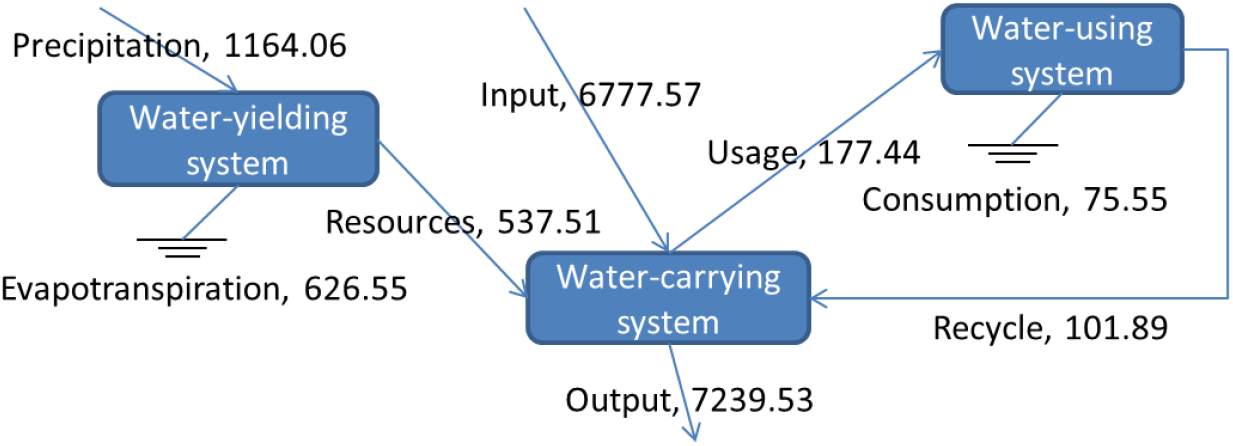
Schematic diagram of the flow network components (Jing MSR: Jingjiang main stream region) of the integrated water resource system in the Yangtze watershed meta-ecosystem (WME) (2014)

### 3.3 Features of the geomorphic topological and spatial structures

Every node in the flow networks of the geomorphic topological structure and geomorphic spatial structure has three elements: an assignment, an input and an output. The input and output describe the flows. The assignment indicates the source, and in these two flow networks, these values were 1 and the size of the corresponding catchment area. The holistic properties (TST, C, H, A, AMI, R, and SR) were determined by ENA (Table 1). Additionally, AMI/H reflects the relative organization degree (ROD).

**Table 1.**
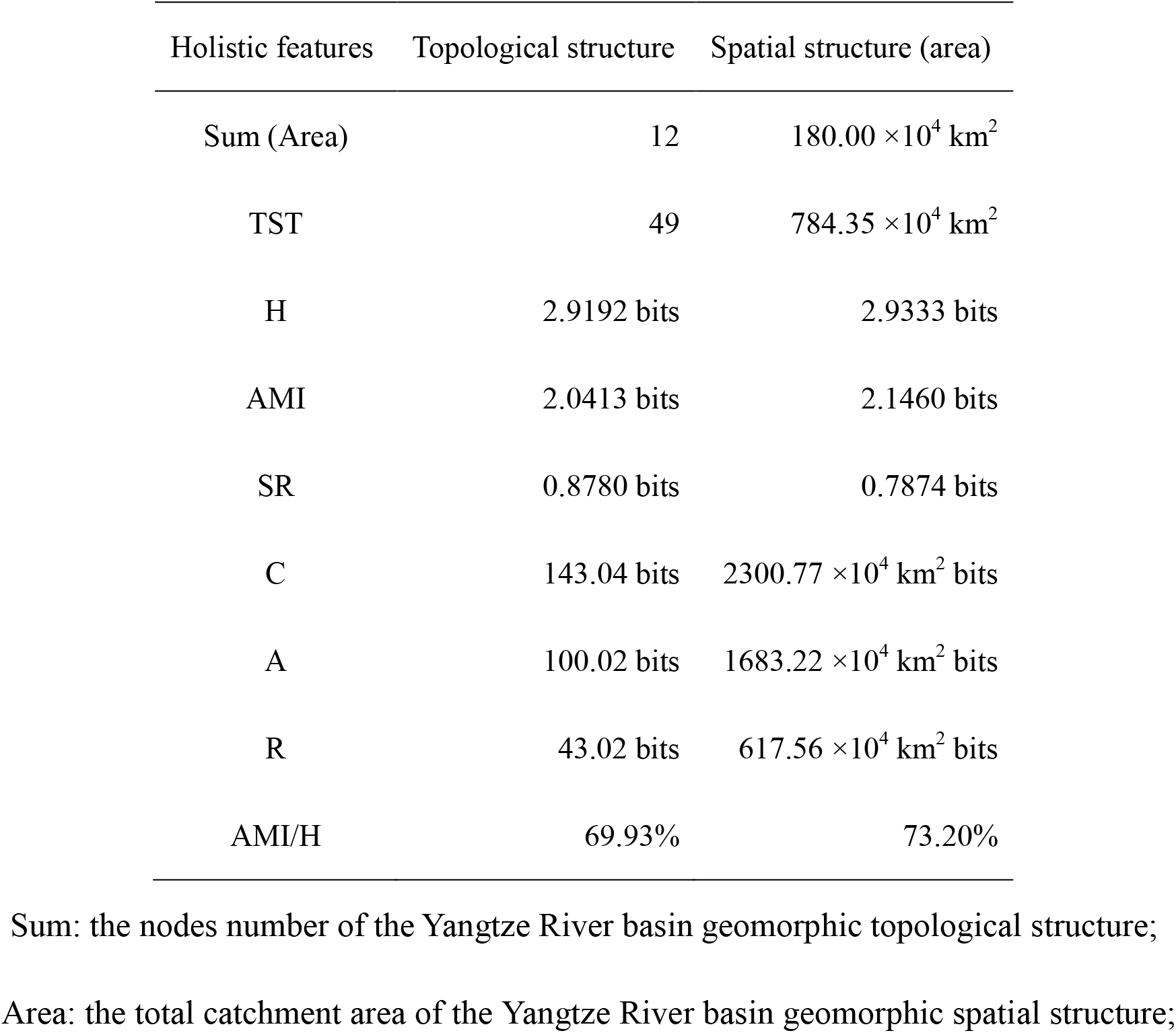
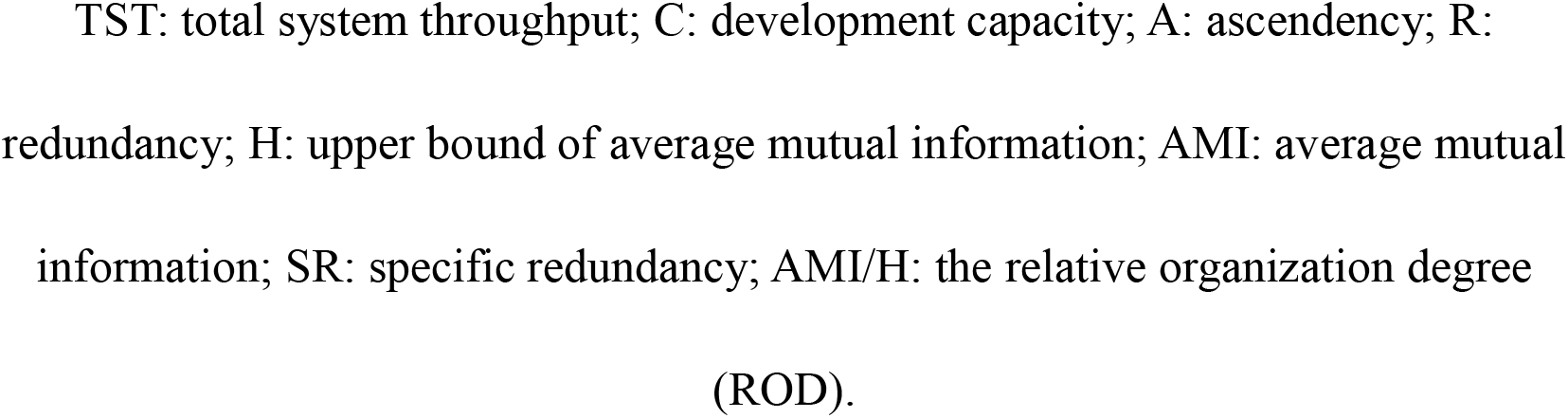
Holistic properties of the geomorphic topological structure and the geomorphic spatial structure (area) of the Yangtze watershed meta-ecosystem (WME)

| Holistic features | Topological structure | Spatial structure (area) |
| --- | --- | --- |
| Sum (Area) | 12 | 180.00 $\times 10^4$ km<sup>2</sup> |
| TST | 49 | 784.35 $\times 10^4$ km<sup>2</sup> |
| H | 2.9192 bits | 2.9333 bits |
| AMI | 2.0413 bits | 2.1460 bits |
| SR | 0.8780 bits | 0.7874 bits |
| C | 143.04 bits | 2300.77 $\times 10^4$ km<sup>2</sup> bits |
| A | 100.02 bits | 1683.22 $\times 10^4$ km<sup>2</sup> bits |
| R | 43.02 bits | 617.56 $\times 10^4$ km<sup>2</sup> bits |
| AMI/H | 69.93% | 73.20% |

Sum: the nodes number of the Yangtze River basin geomorphic topological structure;
Area: the total catchment area of the Yangtze River basin geomorphic spatial structure;
TST: total system throughput; C: development capacity; A: ascendancy; R: redundancy; H: upper bound of average mutual information; AMI: average mutual information; SR: specific redundancy; AMI/H: the relative organization degree (ROD).

### 3.4 Features of the natural water resource system

Based on the data sets (2006-2015) of the Changjiang & Southwest Rivers Water Resources Bulletin, the variations in the quantities of precipitation and water resources from 2006 to 2015 are shown, and the holistic properties (TST, C, H, A, AMI, R, and SR) of the natural water resource system of the Yangtze River basin (2006-2015) were identified (Figure 10).

**Figure 10.**
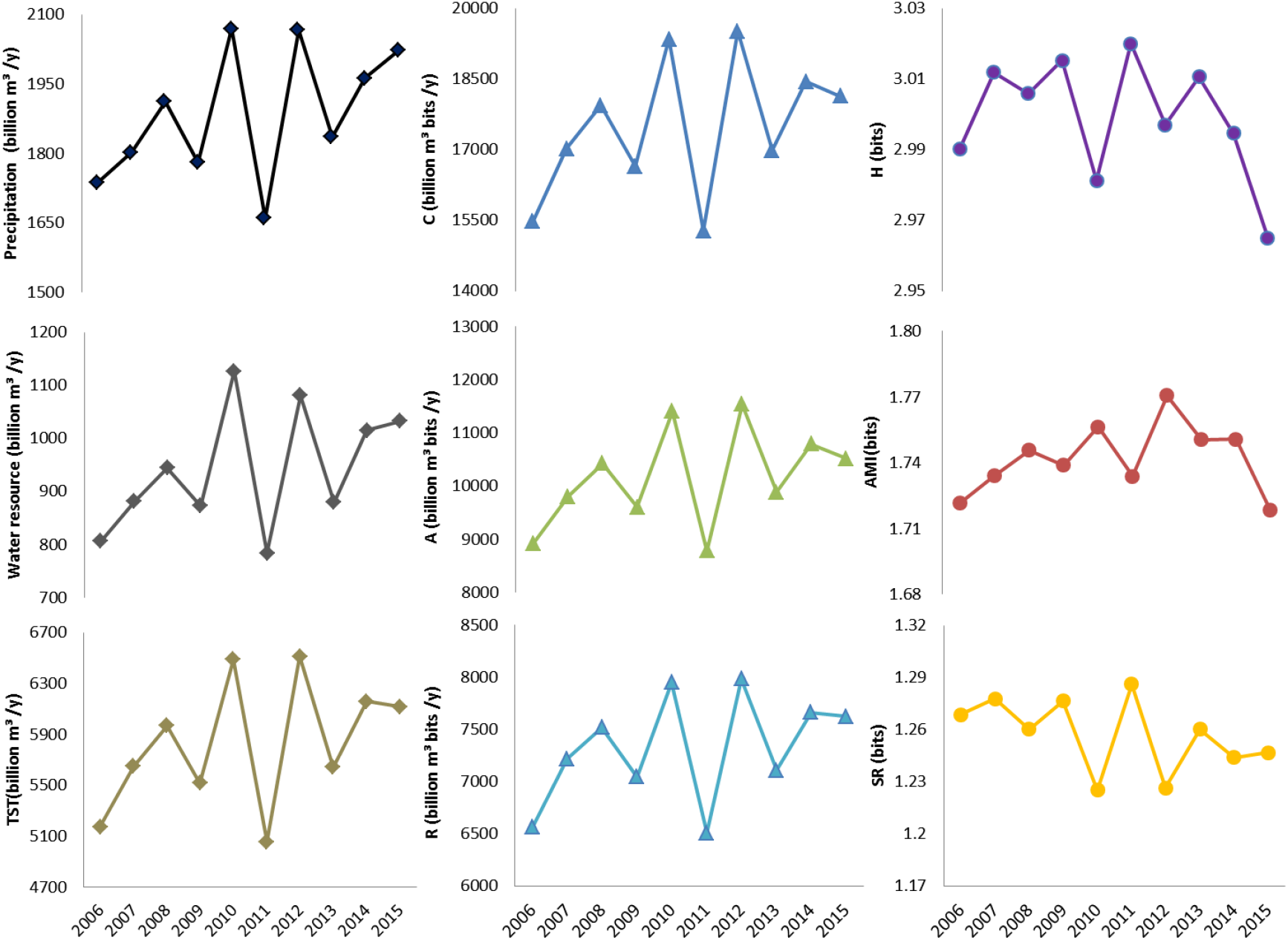
Time series of precipitation, water resources and holistic properties (TST, C, A, R, H, AMI, and SR) of the natural water resource system of the Yangtze River basin (2006-2015). TST: total system throughput; C: development capacity; A: ascendency; R: redundancy; H: upper bound of average mutual information; AMI: average mutual information; SR: specific redundancy.

### 3.5 Features of the integrated water resource system

Based on the data sets (2010-2015) of the Changjiang & Southwest Rivers Water Resources Bulletin, the variations in the TST of the integrated water resource system from 2010 to 2015 is shown, and the holistic properties (TST, C, H, A, AMI, R, and SR) of the integrated water resource system of the Yangtze River basin (2010-2015) were identified (Figure 11).

**Figure 11.**
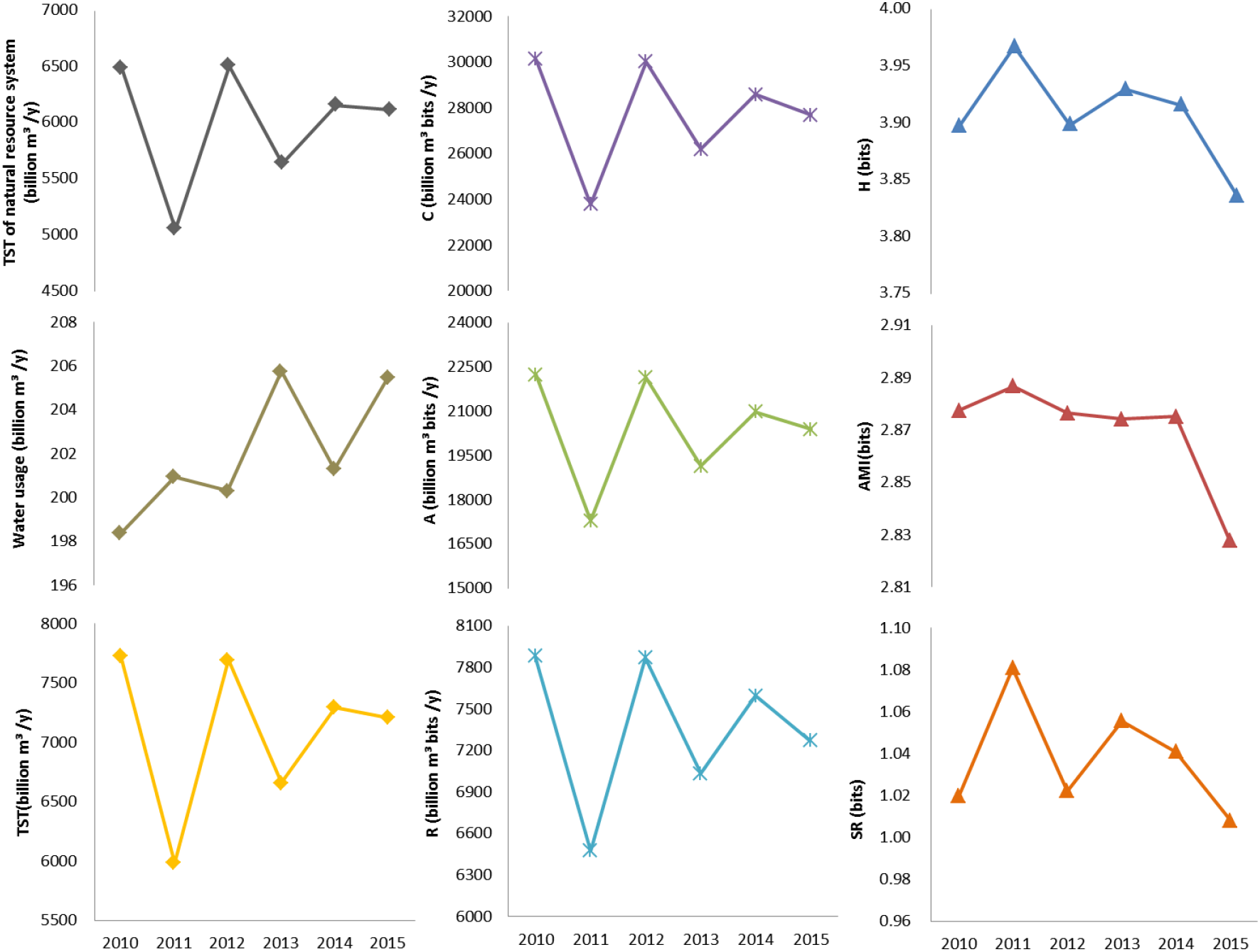
Time series of the TST of the natural water resource system, sum of water usage, and time series of holistic properties (TST, C, A, R, H, AMI, and SR) of the integrated water resource system of the Yangtze River basin (2010-2015). TST: total system throughput; C: development capacity; A: ascendency; R: redundancy; H: upper bound of average mutual information; AMI: average mutual information; SR: specific redundancy.

## 4 Discussion

### 4.1 Different WME construction

The flow network of the geomorphic topological structure is the basic flow network of the geomorphic spatial structure based on the corresponding topological structure. In the Yangtze WME, the sum, TST, AMI, and H of the flow network of the geomorphic topological structure were 12, 49, 2.0413 bits and 2.9192 bits, respectively. The normalized flow network of the geomorphic spatial structure (its sum was 12) had larger TST (activity, 52.29), AMI (organization, 2.1460 bits), and H (upper bound of AMI, 2.9333 bits) values. This finding suggests that the information provided by the geomorphic spatial structure increased the activity and organization of the WME. Similarly, the flow network of the geomorphic spatial structure is the basic flow network of the IWE based on the corresponding spatial structure.

Because there are (1) different minimum subwatershed areas and (2) various strategies for assigning stream order, the construction of a WME can vary. Different methods of WME construction result in different flow networks and geomorphic topological and spatial structures. Then, different IWE flow networks are created based on this WME, and this process affects the holistic properties of these flow networks. For example, if the Yangtze WME is reconstructed following the main-branch strategy, then its geomorphic topological and spatial structure flow networks (Figure 12) are different from the previous ones (Figure 7). Obviously, these changes would affect all the flow networks of the Yangtze IWE based on this WME, as well as the holistic properties of these flow networks.

**Figure 12.**
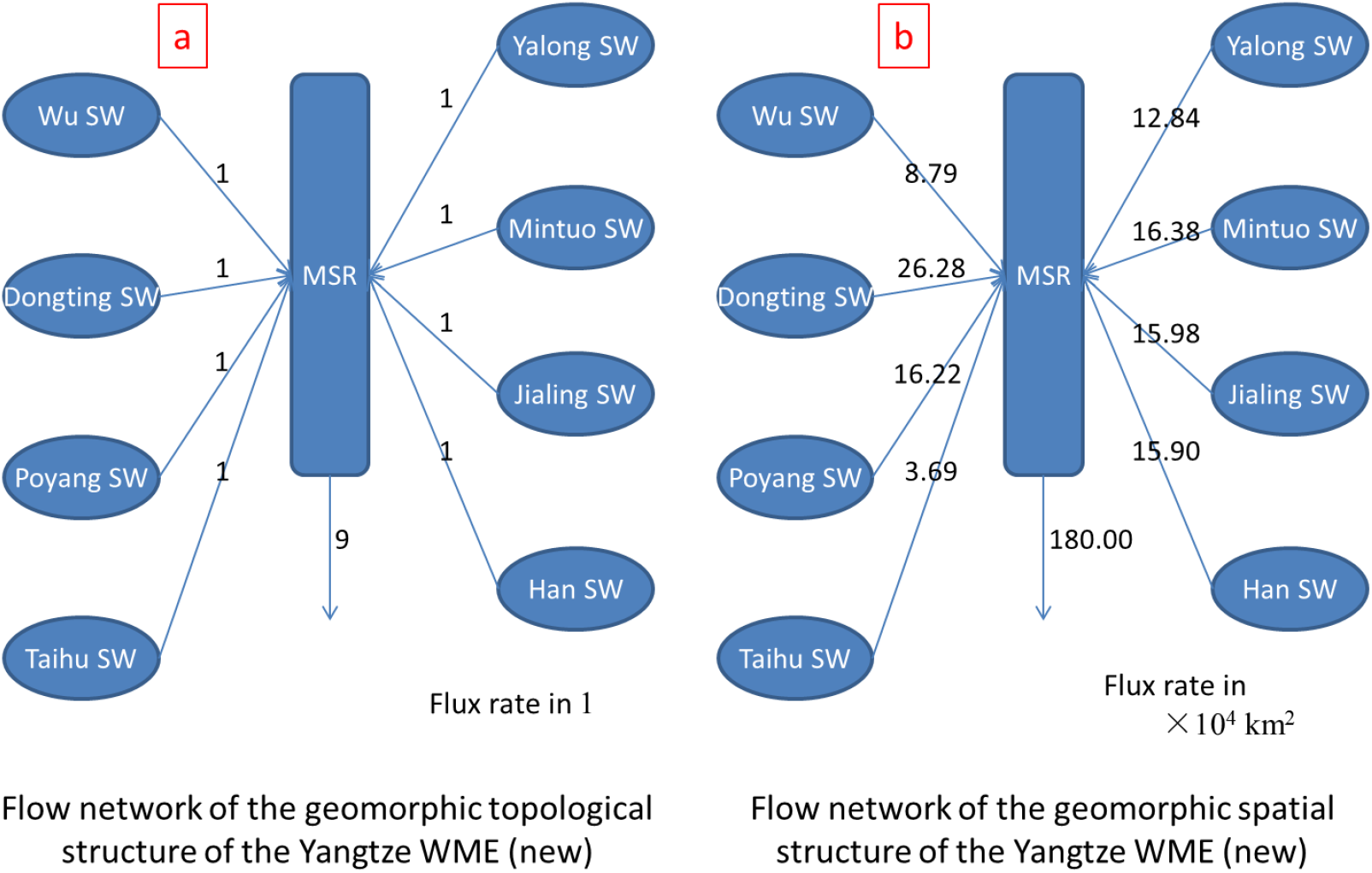
Schematic diagrams of the flow networks of the geomorphic topological structure (a) and geomorphic spatial structure (b) of the new Yangtze watershed meta-ecosystems (WME) constructed based on the main-branch strategy. SW: subwatershed and MSR: main stream region.

### 4.2 Development of the geomorphologic structure of the watershed

The development of the geomorphic structure of the watershed has two elements: (1) extending, which expands a watershed via headward erosion and river capture, and (2) reorganizing, which divides subwatersheds within a watershed due to headward erosion and river capture. In the flow network of the geomorphic spatial structure of the watershed, extending the watershed geomorphic structure means increasing the total area of the watershed, while reorganizing the structure reflects variations in the TST, AMI and A values of the flow network. Of course, when a watershed geomorphic structure extends, it is also reorganized.

Following a defined pathway of constructing a WME, one can construct a set of WMEs and corresponding flow networks for a watershed (or a set of different watersheds) in different developmental stages. By comparing the holistic properties of the flow networks of a watershed in different developmental stages, one could perform a longitudinal study of a watershed development. Additionally, by comparing the holistic properties of the flow networks of a set of different watersheds, one could cluster these watersheds and show the developmental state of each watershed.

For example, based on the evolutionary pattern of the Yangtze River basin described by river captures (Clark et al., 2004), a set of WMEs and corresponding flow networks could be constructed (Figure 13); then, their holistic properties could be calculated (Figure 14). Obviously, when a watershed extends, its main holistic properties increase (from Figure 14a to Figure 14f). Moreover, when a watershed reorganizes, its holistic properties may reflect higher organization (from Figure 14f to Figure 14 g). These processes follow the general trend of system evolution proposed by Ulanowicz (1986).

**Figure 13.**
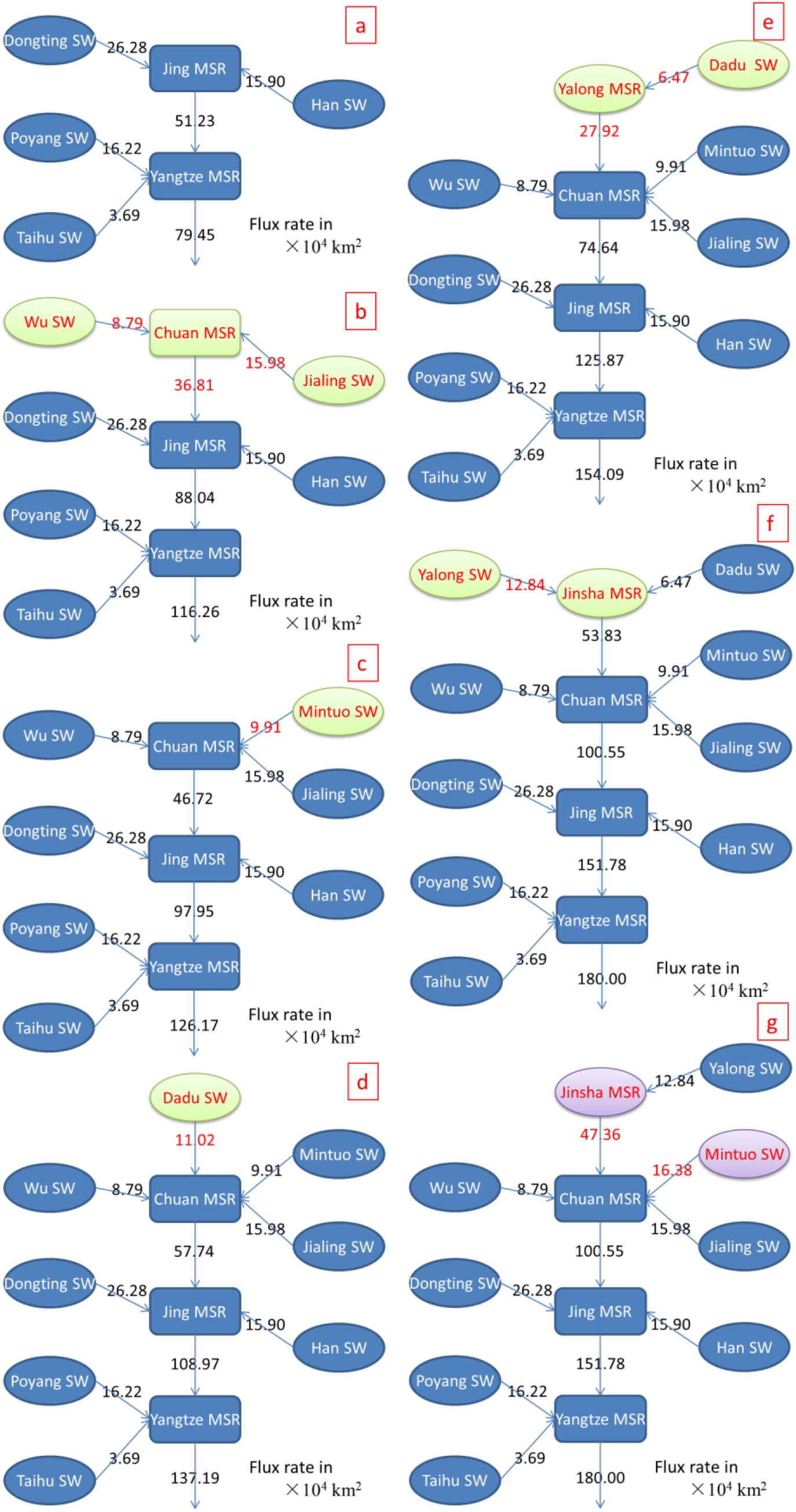
Schematic diagrams of the flow network of the geomorphic spatial structure of the Yangtze watershed meta-ecosystem in different developmental stages (Clark and Schoenbohm et al., 2004). (a) Interpreted original drainage basin of the Lower Yangtze River, where the headwaters of this basin are in the Three Gorges region. (b) Reversal of the Middle Yangtze is initiated by the reversal of a small segment of the river along the main course and the capture of the Jialing River to the east in the Lower Yangtze River basin. (c) Reversal progresses from east to west with the capture of the Min River, (d) the Dadu/Anning River, (e) the Yalong River, (f) and the Jinsha River, and (g) terminates with the capture of the Dadu/Anning River by the Min River, which results in the modern drainage basin morphology. SW: subwatershed and MSR: main stream region.

**Figure 14.**
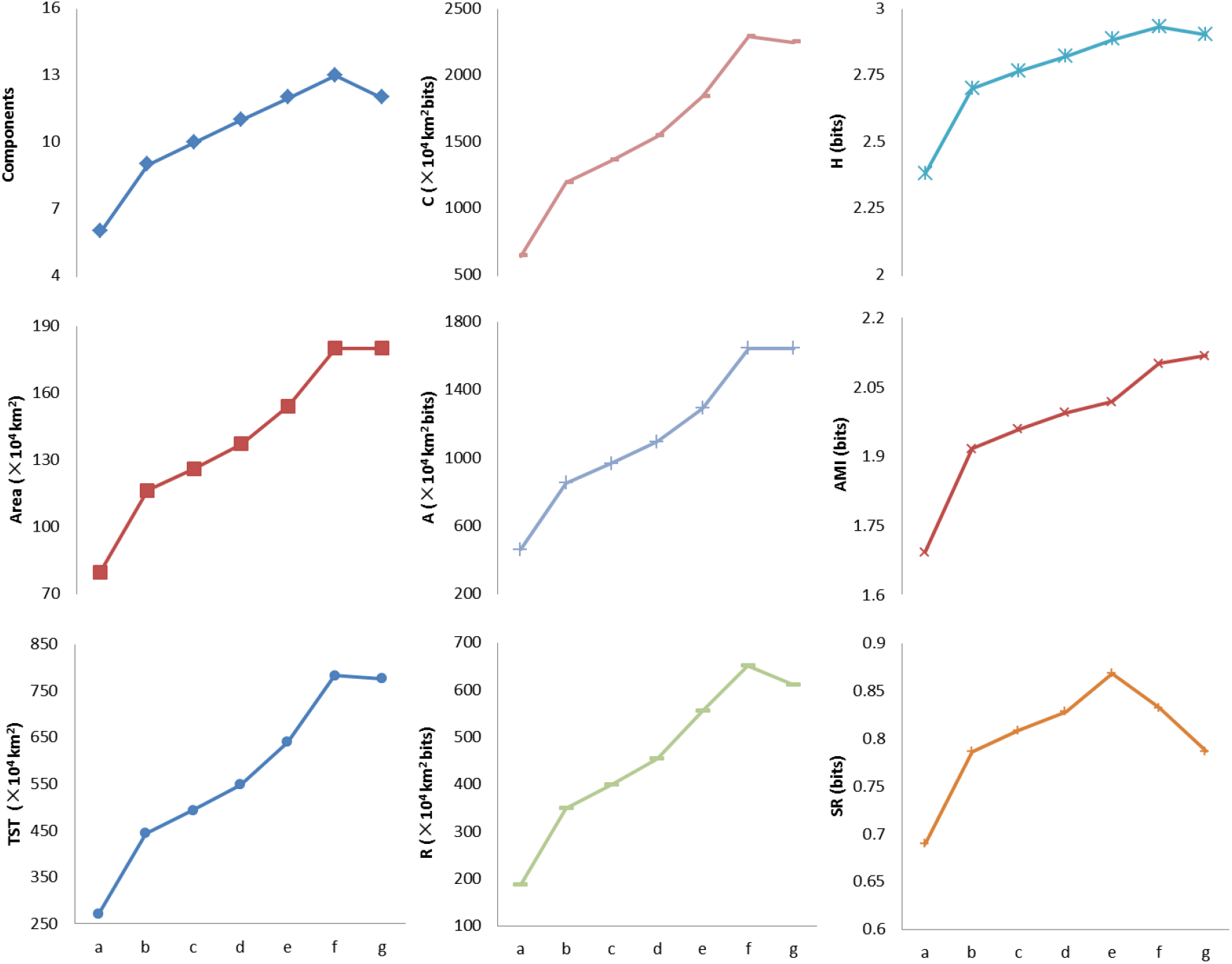
Time series of number of components, watershed area, and holistic properties (TST, C, A, R, H, AMI, and SR) of the flow network of the geomorphic spatial structure of the Yangtze watershed meta-ecosystem in different developmental stages (Figure 13). TST: total system throughput; C: development capacity; A: ascendency; R: redundancy; H: upper bound of average mutual information; AMI: average mutual information; SR: specific redundancy.

### 4.3 Variations in the holistic properties of the water resource system

In the natural water resource system, the variations in TST (activity) were similar to those in precipitation and water resources, and C, A, and R exhibited the same variations as TST, although H, AMI and SR exhibited different fluctuations as mentioned in the Section 3.4. These observations suggest that the variations in C, A, and R generally depend on variations in TST, and fluctuations in TST are mainly due to variations in precipitation and water resources. This finding suggests that the spatial patterns of precipitation and water resources in the Yangtze River basin did not significantly from 2006-2015. This result shows that the annual precipitation trend from 2006-2015 is the continuation of the trend from 1995-2011 (Chen et al., 2014). However, the fluctuations in AMI and SR suggest that the natural water resource system may have experienced high spatial circulation efficiency and low resilience, especially from 2006 to 2012.

In the integrated water resource system, the variation in TST was similar to that in the TST of the natural water resource system, although water usage exhibited a different fluctuation pattern. Additionally, C, A and R exhibited the same fluctuation pattern as TST as mentioned in the Section 3.5. This result suggests that the fluctuation patterns of C, A, and R generally depended on the variations in TST, and the variations in TST were mainly due to fluctuations in the TST of the natural water resource system. Therefore, water usage did not significantly change the spatial pattern of the flow network of the integrated water resource system in the Yangtze River basin between 2010 and 2015, although the sum of water use gradually increased over this period. However, the fluctuations in H, AMI and SR suggest that the integrated water resource system experienced lower and lower spatial circulation efficiency and resilience from 2010 to 2015.

### 4.4 Inner cycles and system resilience

Bidirectional flows exist between subwatersheds and MSRs in the flow network of a natural water resource system (especially in ecological flow networks and socio-economic flow networks), and the bidirectional flows (inner cycles) increase the holistic resilience of the flow network of the natural water resource system. For example, in the flow network of the natural water resource system of the Yangtze WME, R was higher when the inner cycles were taken into account, although the A was nearly invariable (Figure 15).

**Figure 15.**
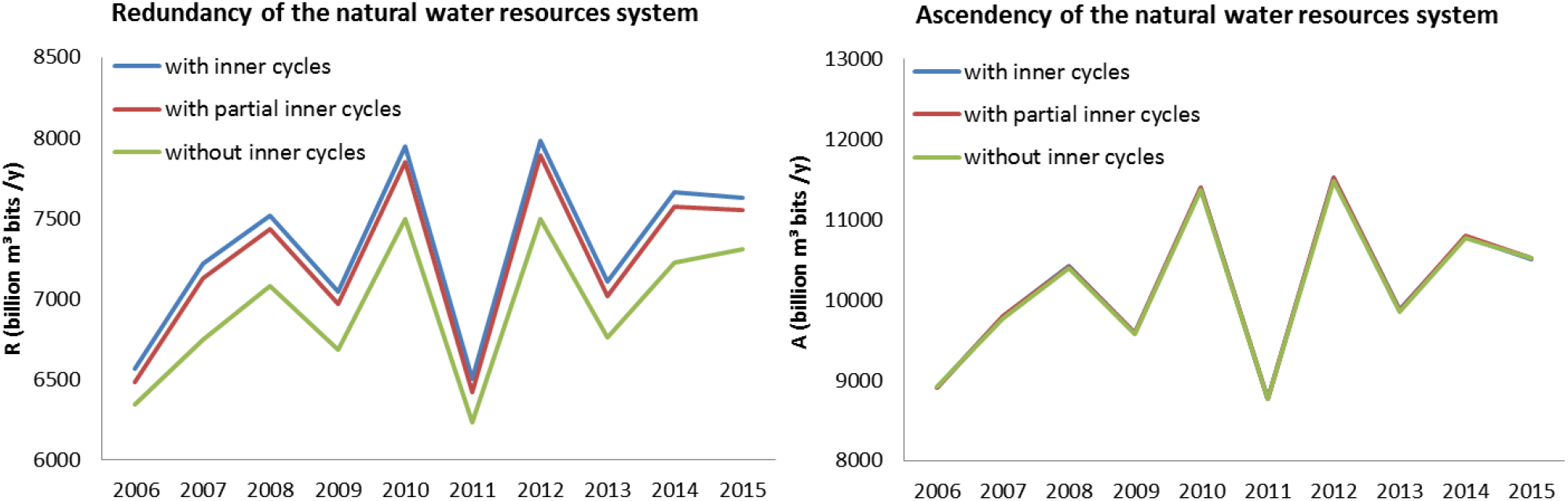
Time series of redundancy (R) and ascendency (A) of the natural water resource system of the Yangtze River basin (2006-2015) under three scenarios. The flow network with inner cycles is the flow network including water flows (1) from the main stream region to the Dongting lake subwatershed and Poyang Lake subwatershed and the water flows (2) between the Taihu Lake subwatershed and the main stream region/ Qiantang watershed. The flow network with partial inner cycles is the flow network including only the water flows listed in (1) (above). The flow network without inner cycles is the flow network without water flows (1) and (2).

Considering the variations in the annual water discharge of the Yangtze River (Figure 16a), the continuous declining trends of flow diversion at the three outlets along the Jingjiang reach (Figure 16b) and the fluctuating by declining trend in the reflow volume from Yangtze River to Poyang Lake (Figure 16c) over the past 60 years, the system resilience of the natural water resource system of the Yangtze River basin has decreased over the past 60 years, although the volume of water transferred from the MSRs of the Yangtze River basin and Qiantang River basin to the Taihu Lake subwatershed increased over the same period (Figure 16d).

**Figure 16.**
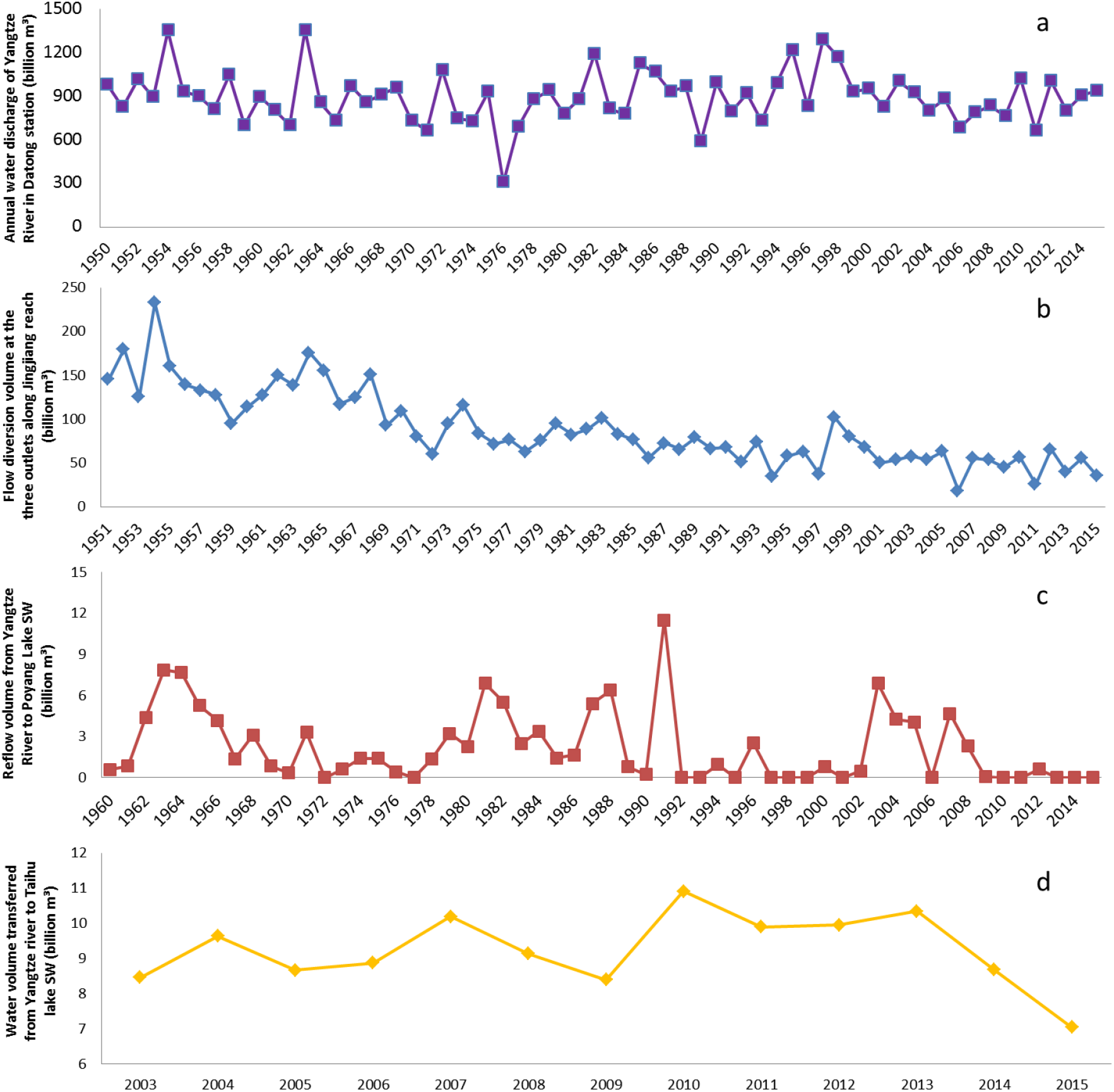
Annual water discharge time series from Datong station in the Yangtze River basin (a). Flow diversion volume time series from the Yangtze River to the Dongting Lake subwatershed at the three outlets along the Jingjiang reach (b). Reflow volume time series from the Yangtze River to the Poyang Lake subwatershed at Hukou (c). Time series of the water volume transferred from the MSRs of the Yangtze River basin and Qiantang River basin to the Taihu Lake subwatershed (d).

### 4.5 Spatial properties of precipitation and eco-hydrological conditions

By comparing the holistic properties of the watershed geomorphic spatial structure with the holistic properties of the watershed natural water resource system without inner cycles, the holistic spatial properties of precipitation and eco-hydrological conditions can be determined. For example, A/C (or AMI/H) reflects the ROD of a WME, and the ROD of the watershed geomorphic spatial structure can be compared with the ROD of the watershed natural water resource system without inner cycles to determine the ROD of spatial precipitation and eco-hydrological conditions in the watershed.

For example, the ROD of the watershed geomorphic spatial structure in the Yangtze River basin is 73.20%, and the ROD of the natural water resource system without inner cycles in the Yangtze River basin varies from 58.42%~60.51%. Thus, the ROD of spatial precipitation and eco-hydrological conditions is 0.7981~0.8266, which suggests that the spatial properties of precipitation and eco-hydrological conditions have a negative effect on the natural water resource system of the watershed (Figure 17).

**Figure 17.**
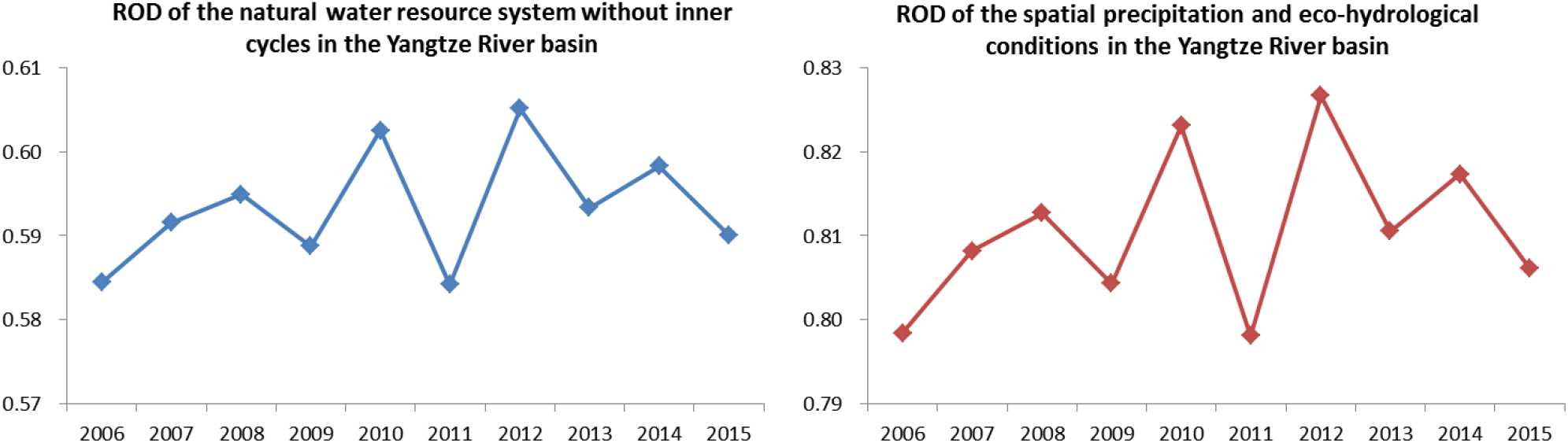
Time series of the relative organization degree (ROD) values of the natural water resource system without inner cycles in the Yangtze River basin and the spatial precipitation and eco-hydrological conditions in the Yangtze River basin

### 4.6 ENA of the subwatershed meta-ecosystem

QDFIWE is a general framework that can be used to identify the holistic properties of every watershed or subwatershed in the hierarchical watershed-subwatershed system. For example, the Yangtze River basin was analyzed as the WME, but the subwatersheds of the Yangtze River basin, such as the Poyang Lake basin, could also be analyzed as WMEs. Following the watershed resource divisions of the Jiangxi Water Resources Bulletin (http://www.jxsl.gov.cn/slgb/szygb/index.html), the Poyang Lake WME was constructed (Figure 18), and the holistic properties of the natural water resource system were identified (Figure 19).

**Figure 18.**
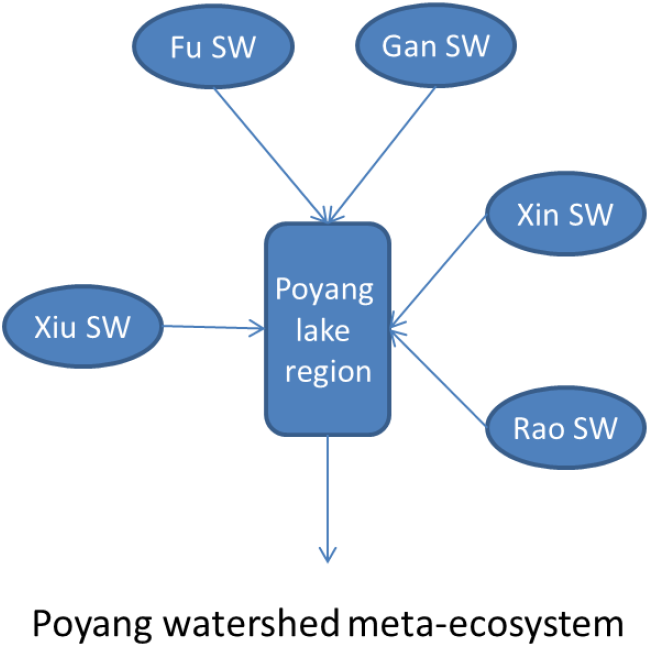
Schematic diagram of the Poyang watershed meta-ecosystem (WME) constructed based on the watershed resource divisions of the Jiangxi Water Resources Bulletin (http://www.jxsl.gov.cn/slgb/szygb/index.html). SW: subwatershed and MSR: main stream region.

**Figure 19.**
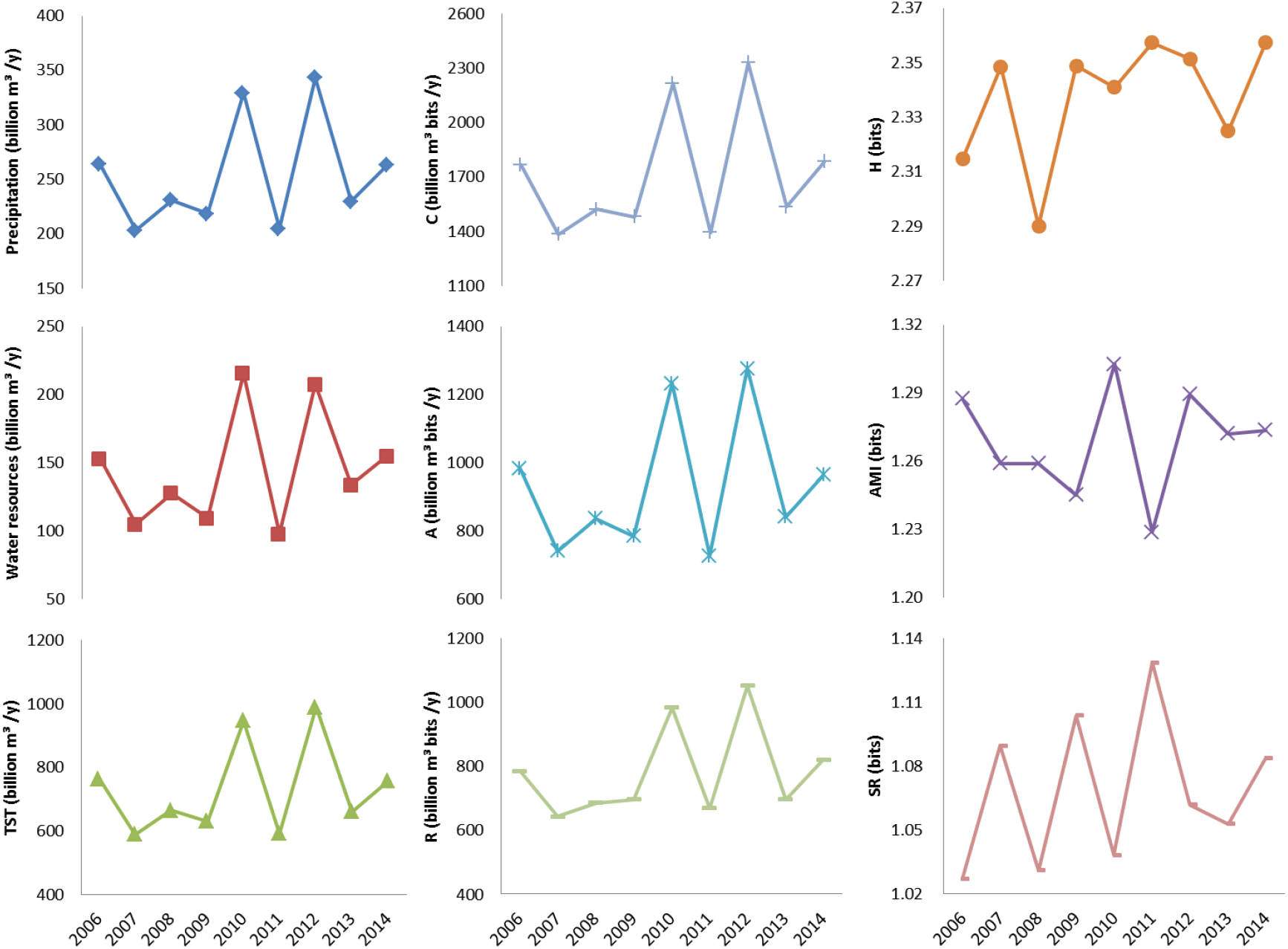
Time series of precipitation, water resources and the holistic properties (TST, C, A, R, H, AMI, and SR) of the natural water resource system of Poyang Lake basin (2006-2014). TST: total system throughput; C: development capacity; A: ascendency; R: redundancy; H: upper bound of average mutual information; AMI: average mutual information; SR: specific redundancy.

In the natural water resource system of Poyang Lake basin, the variations in C, A, and R generally depend on flucations in TST, and the fluctuations in TST are mainly due to variations in precipitation and water resources. Thus, the spatial properties of the precipitation and eco-hydrological conditions in Poyang Lake basin did not significantly change from 2006 to 2014. However, the fluctuations in AMI and SR suggest that the natural water resource system experienced lower and lower spatial circulation efficiency and higher and higher resilience from 2006 to 2014.

Subwatersheds must be prioritized for better management of the overall watershed, and many approaches have been developed to prioritize subwatersheds (Jain and Das, 2010; Chowdary et al., 2013; Jaiswal et al., 2014; Adhami and Sadeghi, 2016). In the QDFIWE, the resilience of watersheds (or subwatersheds) provides a useful and important indicator for prioritizing a set of watersheds (or subwatersheds) and improving watershed management at various subsystem levels of IWEs and the overall IWE level.

## 5 Conclusion

In this study, we introduced the QDFIWE, which provides a general method of quantitatively describing the geomorphic subsystem, hydrologic subsystem, ecological subsystem and socio-economic subsystem of an IWE simultaneously.

1. In the QDFIWE, based on different settings of the minimum subwatershed area and various strategies of assigning stream order, one can construct different WMEs. These different WMEs can satisfy the different requirements of IWE analysis.
2. Following the QDFIWE, one can generate time series of the holistic properties of a watershed based on its geomorphic features, hydrologic features, ecological features and socio-economic features. These time series of holistic properties reflect the geomorphic evolution, hydrologic evolution, ecological evolution and socio-economic evolution of the IWE.
3. Based on the construction of WMEs and flow networks using similar strategies, any number of IWEs can be analyzed, compared, classified and clustered by identifying their holistic properties.

The application of this method to the Yangtze River basin revealed that:

1. When the geomorphic structure of the watershed was extended, its main holistic properties increased too. Additionally, when the watershed geomorphic structure was reorganized without extension or shrinkage, its holistic properties tended toward higher organization. These results reflect the general propensity of system evolution.
2. As the inner cycles in the natural water resource system of the Yangtze River basin decreased over the past 60 years, the resilience of the natural water resource system of the Yangtze River basin decreased too. Thus, increasing the inner cycles will benefit the resilience of flow networks constructed based on the hydrologic, ecological and socio-economic spatial processes of IWE.
3. According to the ROD, A/C or AMI/H of the spatial precipitation and eco-hydrological conditions and the holistic properties of the geomorphic spatial structure and natural water resource system without inner cycles, we found that the spatial properties of precipitation and eco-hydrological conditions had a negative effect on the natural water resource system of the Yangtze IWE. Thus, more indicators could be determined or created based on the holistic properties (TST, C, H, A, AMI, R, and SR) of the IWE to provide additional information.

## Acknowledgments

This research did not receive any specific grant from funding agencies in the public, commercial, or not-for-profit sectors.

